# Modeling Prenatal Immune Activation in Human Brain Organoids Uncovers IL-6–Dependent Interneuron Dysmaturation

**DOI:** 10.64898/2026.05.26.728058

**Authors:** Ava V. Papetti, Ziyuan Ma, Michelle Ng, Mengmeng Jin, Steven W. Levison, Peng Jiang

**Affiliations:** Department of Cell Biology and Neuroscience, Rutgers University-New Brunswick, Piscataway, NJ 08854, USA; Department of Pharmacology, Physiology & Neuroscience, New Jersey Medical School, Rutgers University, Newark, NJ, 07103, USA

## Abstract

Prenatal inflammation has been associated with an increased likelihood of the child developing neurodevelopmental conditions, such as autism spectrum disorder (ASD). Several pro-inflammatory cytokines are significantly upregulated and play critical roles during immune activation, with interleukin-6 (IL-6) being particularly prominent. However, the specific impact of elevated IL-6 levels on human neural development remains to be elucidated. To address this, we established a human pluripotent stem cell-based forebrain organoid model enriched for multiple interneuron lineages, validated through immunohistochemistry and transcriptomic alignment with human fetal reference datasets. We showed IL-6 responsiveness via the activation of the JAK/STAT pathway and characterized downstream effects using bulk and single-nucleus RNA sequencing (RNA-seq). Bulk RNA-seq at the end of IL-6 exposure uncovered activation of inflammatory pathways, upregulation of MHC-I machinery, and early disruption of GABAergic signaling programs. Notably, single-nucleus RNA-sequencing performed one month after IL-6 withdrawal revealed a persistent inflammatory transcriptional signature across interneuron development, accompanied by accelerated progression through maturation stages and altered interneuron subtype output. Together, these findings demonstrate that transient prenatal IL-6 exposure is sufficient to reshape human interneuron fate specification and maturation trajectories, providing mechanistic insights into neuroimmune contributions to ASD.

## Introduction

Epidemiological evidence has pinpointed several environmental risk factors during pregnancy, ranging from maternal infection, autoimmune conditions, allergies, and air pollution, that are associated with an autism spectrum disorder (ASD) diagnosis later in the child’s development^1–3^. A shared mechanism downstream from these exposures is immune system activation. However, the precise mechanisms through which inflammatory signaling exerts cell-type-specific effects on neuronal development and maturation—and thereby promotes ASD symptomology—remain elusive.

The immune response is mediated by pro-inflammatory cytokines and chemokines, some of which can cross the placenta and the blood-brain barrier in the developing fetus^4^. These pro-inflammatory cytokines can then guide changes in brain development linked to the onset of neurodevelopmental disorders. One cytokine that can cross both the placenta and the fetal blood-brain barrier^5,6^ and has consistently been shown to be associated with the manifestation of ASD-like features in rodent models and in humans is interleukin-6 (IL-6). A seminal murine study found that IL-6 injection alone into pregnant dams at E12.5 produced significant gene expression changes within the frontal cortex, followed by social behavioral deficits in the adult offspring. Remarkably, these changes were prevented when IL-6 antibodies were introduced midgestation^7^. Moreover, IL-6 receptor knockout mice failed to develop the social interaction deficits observed in wild-type mice upon prenatal administration of a viral mimetic^7^, demonstrating that IL-6 is both necessary and sufficient to induce ASD phenotypes. Human studies have correlated maternal levels of IL-6 during pregnancy with altered functional brain connectivity in infants aged 18-24 months, changes in amygdala volume and connectivity at 24 months, and changes in white matter composition^8–10^. Furthermore, IL-6 has been observed to be elevated in fetal circulation and cerebrospinal fluid^4,11^, as well as in serum collected from children at ages 2-4 years old who were later diagnosed with ASD^12^. Increased levels of IL-6 have also been observed to be associated with stereotypical behaviors in ASD^13^, emphasizing the connection between elevated IL-6 to brain structure and organization and behavior. Taken together, these data point to IL-6 as a critical mediator of neuroanatomical, molecular, and behavioral changes that contribute to ASD pathogenesis.

Several studies using rodent models to uncover the effects of maternal immune activation (MIA) on neurodevelopment have converged on alterations in the subtype-specific abundance and activity of cortical GABAergic interneurons^14–18^. Interneurons have been reported to comprise up to 50% of neurons in the human cortex^19,20^, and they play a critical role in maintaining the excitation-inhibition (E/I) balance within neural circuits^21^. A prevailing framework for ASD pathology implicates the dysregulation of E/I balance, which is dependent on proper interneuron maturation and functional integration^22–24^.

While studies using rodent models have highlighted the vulnerability of interneuron lineage cells to neuroinflammation, the translation of these immune-mediated changes to humans remains largely unknown. Further constraining our understanding of early etiological mechanisms is the inaccessibility of prenatal human brain tissue, a developmental window that precedes the clinical diagnosis of ASD. Moreover, human interneurons develop from distinct progenitor pools with unique transcriptional profiles that give rise to over a dozen diverse subclasses^20,25–30^, underscoring the need for a model system that captures the multifaceted aspects of their development.

Human pluripotent stem cell (hPSC)-derived forebrain organoid models, which can reproduce human-specific interneuron diversity, provide a platform for studying the effects of immune challenges on stages of fetal brain development that have been inaccessible. This is particularly relevant for modeling ASD, which, although diagnosed during early childhood, has prenatal origins. A recent study exposed dorsal forebrain organoids to IL-6 and showed modest changes in neural progenitor cell (NPC) differentiation toward excitatory neurons^31^. However, the model in this study lacked ventral forebrain NPCs, which give rise to the majority of cortical interneurons. Although the analysis focused on excitatory neurons, differential gene expression analyses (DEG) nonetheless revealed changes in genes associated with interneuron development following IL-6 treatment^31^. In the present study, we established a human induced pluripotent stem cell (hiPSC)–based organoid model enriched for interneuron progenitors to more precisely investigate the effects of IL-6 exposure on interneuron development. This new system enables simultaneous assessment of IL-6 effects on lineages originating from multiple subdomains of the forebrain. By introducing IL-6 to organoids early after inducing neuronal differentiation, we have characterized how IL-6 alters the trajectory of human interneuron development. Our analyses reveal sustained inflammatory signatures spanning early progenitor stages through the acquisition of mature interneuron fate, along with changes in maturation dynamics and subtype lineage specification.

## Methods

### Human iPS cell lines, generation, and culture

Three healthy control male hiPSC lines were used in this study. All lines were fully characterized by karyotyping, gene expression, and PluriTest analysis^103^, as described in our previous work^26^. All hiPSCs were cultured on dishes coated with hESC-qualified Matrigel (Corning) and maintained in mTeSR Plus medium (STEMCELL Technologies) under a feeder-free condition. hPSCs were passaged with ReLeSR media (STEMCELL Technologies) once per week.

### Differentiation and culture of dorsoventrally patterned NPCs

Human NPCs were generated from all three hPSC cell lines using a previously established protocol^26,104^. Briefly, expanded pNPCs were dissociated into single cells using TrypLE Express (Thermo Fisher Scientific) and seeded at a density of two million cells per well in suspension in ultra-low attachment 6-well plates, with 10 µM ROCK inhibitor Y-27632 (Tocris) added on the first day to promote survival. Beginning on the second day, pNPCs were transitioned to a ventralization medium comprised of a 1:1 mixture of Neurobasal (Thermo Fisher Scientific), DMEM/F12 (HyClone), and GlutaMax (Gibco), supplemented with 1x N2 (Thermo Fisher Scientific), 1x B27-RA (Thermo Fisher Scientific), and recombinant human FGF2 (20 ng/ml, Peprotech). To stimulate ventralization, 1 µM Purmorphamine (Cayman Chem) was added to the medium, which was replenished daily for one week. After one week of patterning, spheroids were dissociated into single cells using TrypLE Express and replated onto Growth Factor Reduced Matrigel-coated dishes for adherent expansion. These NPCs were maintained in the same medium without Purmorphamine (NPC Medium) without further morphogen stimulation.

### Brain organoid culture

Brain organoids were generated by seeding a total of 10,000 dvNPCs per well of ultra-low-attachment 96-well plates in the presence of 10 µM ROCK inhibitor Y-27632. Organoids were cultured in NPC Medium for four days. On Day 4, the organoids were transferred to ultra-low-attachment 6-well plates (6-8 organoids per well) and cultured with Glial Progenitor Cell medium composed of DMEM/F12, supplemented with 1x N2, FGF2 (10 ng/ml), PDGF-AA (10 ng/ml, Peprotech), IGF1 (10 ng/ml, Peprotech), and dibutyryl-cyclic AMP (1 uM, Sigma) for two weeks (day 18). To promote neural differentiation, organoids were cultured in Differentiation Medium, comprised of a 1:1 mixture of Neurobasal and DMEM/F12, supplemented with 1x N2, BDNF (10 ng/ml, Peprotech), GDNF (10 ng/ml, Peprotech), dibutyryl-cyclic AMP (1 uM), and ascorbic acid (200 nM, Sigma), from day 18 onwards. The medium was replenished every two days, and starting from day 4, the cell culture plates were kept on an orbital shaker at a speed of 85 rpm. Organoids were collected at Day 30 and 60 for experiments. All organoid cultures were maintained at 37°C in a humidified incubator with 5% CO_2_ in atmospheric air.

### IL-6 and sIL-6R treatment

Human recombinant IL-6 (PeproTech) and human recombinant sIL-6R (R&D Systems) were diluted in 0.1% BSA in PBS. On Day 21 the medium was completely replenished with fresh Differentiation Medium with supplemented 8.8 ng/ml (422.6 pM) of IL-6 and 16 ng/ml (422.6 pM) sIL6R. Complete media changes were conducted daily until IL-6 withdrawal on Day 31.

### Immunostaining, image acquisition, and cell quantification

Organoids were fixed with 4% paraformaldehyde for two hours, then cryoprotected by immersion in 25% sucrose solution. Then, organoids were embedded in OCT compound and frozen for cryosectioning. Cryosections of 12-15µm were obtained for immunohistochemistry. For immunostaining, organoids were blocked with a solution containing 5% goat serum in PBS with 0.2% Triton X-100 at room temperature for 1 hour. Primary antibodies were diluted in the blocking solution and incubated with the organoids overnight at 4°C. Following primary antibody incubation, sections were washed with PBS and incubated with secondary antibodies at room temperature for 1 hour. After additional PBS washing, slides were mounted with anti-fade Fluoromount-G medium containing DAPI (1,4,6-diamidino-2-phenylindole dihydrochloride, Southern Biotechnology). Fluorescent images were captured using a Zeiss 800 confocal microscope.

### RNA isolation and quantitative reverse transcription PCR

RNA extraction was performed using the Qiagen Mini kit (Qiagen). Reverse transcription was done using the SuperScript™ IV VILO™ Master Mix (Thermo Fisher Scientific). Real-time PCR was performed on the ABI 7500 Real-Time PCR System using the TaqMan Fast Advanced Master Mix (Thermo Fisher Scientific). The 2^-ΔΔCt^ method was used to calculate relative gene expression after normalization to GAPDH as an internal control.

### Bulk RNA sequencing library preparation and data analysis

Libraries for bulk RNA-seq were prepared using the Illumina TruSeqV2 kit (Illumina, San Diego, CA) following the manufacturer’s protocol. Sequencing was performed on a Novaseq X Plus with 150 bp paired-end reads. The sequenced reads were quality-checked with FastQC (v0.12.1) and trimmed for adaptor and low-quality sequences using Trim Galore (v0.6.10). Trimmed reads were mapped to the GRCh38 reference genome and GENCODE v44 primary assembly using STAR^105^ (v2.7.11a). Read count extraction and normalization were performed with RSEM^106^ (v1.3.1), and differential expression analysis (DEG) was conducted using the edgeR^107^ R package (v4.8.2). DEGs were selected based on the following criteria: |log2 Fold Change| > 0.5 and Benjamini-Hochberg corrected FDR < 0.2 between IL-6-treated and control organoids. Heatmaps were generated using the pheatmap R package (v1.0.12). Differentially expressed genes were visualized using a volcano plot generated with the ggplot2 R package (v4.0.2). Transcription factor activity changes were estimated using the DoRothEA R package (v.1.22.0), and gene set enrichment analysis (GSEA) was performed against MSigDB C2 curated gene sets using the clusterProfiler package (v.4.18.4)

### Single-nucleus preparation and snRNA-seq library construction

Single nuclei were prepared using Nuclei PURE Prep kit (Sigma) following the manufacturer’s protocols. Isolated nuclei suspensions were loaded onto the Chromium Controller for snRNA-seq library preparation using the Chromium Next GEM Single Cell 3′ Kit v3.1 in combination with the Dual Index Kit TT Set A, following the manufacturer’s protocols. Library quality and fragment size distributions were assessed using an Agilent 2100 Bioanalyzer. Sequencing was performed on an Illumina NovaSeq 6000 system using S4 flow cells.

### Single-nucleus RNA sequencing data analysis

Raw sequencing reads were aligned to the human GRCh38 reference genome using Cell Ranger (v7.1.0). Genes expressed in fewer than 3 cells were excluded from downstream analyses. Nuclei were retained if they contained between 200 and 5,000 detected genes and exhibited mitochondrial transcript content below 1%.

Downstream analysis was conducted using the Seurat R package (v5.3.1)^108^. Raw gene count matrices were normalized by regularized negative binomial regression using the *SCTransform()* function (vst.flavor = “v2”), and batch effect was corrected by Canonical Correlation Analysis (CCA)^108–110^. Nonlinear dimensionality reduction was performed using Uniform Manifold Approximation and Projection (UMAP) based on the top 30 principal components. Clustering was performed with a resolution parameter set to 0.4.

Reference postmortem human snRNA-seq datasets were used for cell type correlation and age prediction analyses^42,58^. For the reference dataset, cells were grouped by annotated cell type or age group, and 1,000 cells were randomly sampled from each group. Spearman correlation analysis was performed to compare bulk RNA-seq profiles with pseudobulk profiles generated from single-cell datasets. Cosine similarity analysis was used to compare expression profiles between single-cell datasets.

Volcano plot was produced using EnhancedVolcano R package (v1.28.2) with R v4.5.2. Differentiation potential was inferred using the CytoTRACE^111^ R package (v1.1.0), CytoTRACE scores were computed from normalized expression matrices and used to assess relative differentiation potential across cell populations. Gene ontology (GO) enrichment analysis was performed using the clusterProfiler^112^ R package (v4.18.2). The scProportionTest R package was used to perform permutation tests to assess differences in cluster proportions between experimental groups. Pseudotime trajectories were inferred using the slingshot algorithm^113^.

Gene regulatory networks were inferred using the pySCENIC^114,115^ framework (v0.12.1). Regulon modules were defined based on the Connection Specificity Index (CSI), a context-dependent metric that quantifies the specificity of associations between regulons^114,116^. Briefly, Pearson correlation coefficients (PCCs) were calculated for all pairs of regulon activity scores. For a given pair of regulons, A and B, the CSI was defined as the fraction of regulons whose PCC with both A and B was lower than the PCC observed between A and B. Hierarchical clustering using Euclidean distance was applied to the CSI matrix to identify distinct regulon modules, which were visualized using the ComplexHeatmap^117,118^ R package. Regulon activity scores were calculated using the AddModuleScore() function from the Seurat R package.

### Statistical analysis

All data are represented as means ± SEM. When only two independent groups were compared, significance was determined by using a two-tailed unpaired t-test with Welch’s correction. A p-value of < 0.05 was considered significant. All statistical information, including sample sizes (n) and definitions of statistical significance, is reported in the figure legends. All analyses were performed using GraphPad Prism v.9.

## Results

### Generating Human PSC-Derived Dorsoventrally Patterned Forebrain Organoids to Model Neuroimmune Effects of Interleukin-6

To investigate how pathological levels of IL-6 influence early human neuronal development, we established a hiPSC-derived forebrain organoid system that captures both dorsal and ventral lineage specification. During embryogenesis a sonic hedgehog (SHH) morphogen gradient ventralizes the neuroepithelium, giving rise to 3 discrete structures known as the ganglionic eminences that emerge along the dorsoventral axis. High concentrations of SHH specify medial ganglionic eminence (MGE) specification, while lower concentrations promote lateral ganglionic eminence (LGE) and caudal ganglionic eminence (CGE) identities^32–34^. To generate a mixed progenitor population comprising neuroepithelial cells with both dorsal and ventral forebrain identities, we patterned hiPSC-derived primitive neural progenitor cells (pNPCs) with the SHH pathway agonist Purmorphamine (Pur) for one week to induce ventralization through activation of SHH signaling (Figure 1A). This approach yielded a heterogeneous NPC population expressing markers associated with multiple forebrain domains, including the dorsal forebrain and lateral ganglionic eminence (LGE) markers PAX6 and MEIS2, the medial ganglionic eminence (MGE) marker NKX2.1, the caudal ganglionic eminence (CGE) marker NR2F1, and a smaller subset of OLIG2+ cells, which is expressed in ventral progenitor cells that have the potential to become interneurons^26^ (Figure 1B,C). These data demonstrate that our method captures the diversity of forebrain germinal zones. Importantly, these proportions were reproducible across three independent iPSC lines, indicating robust and consistent patterning.

**Figure 1.**
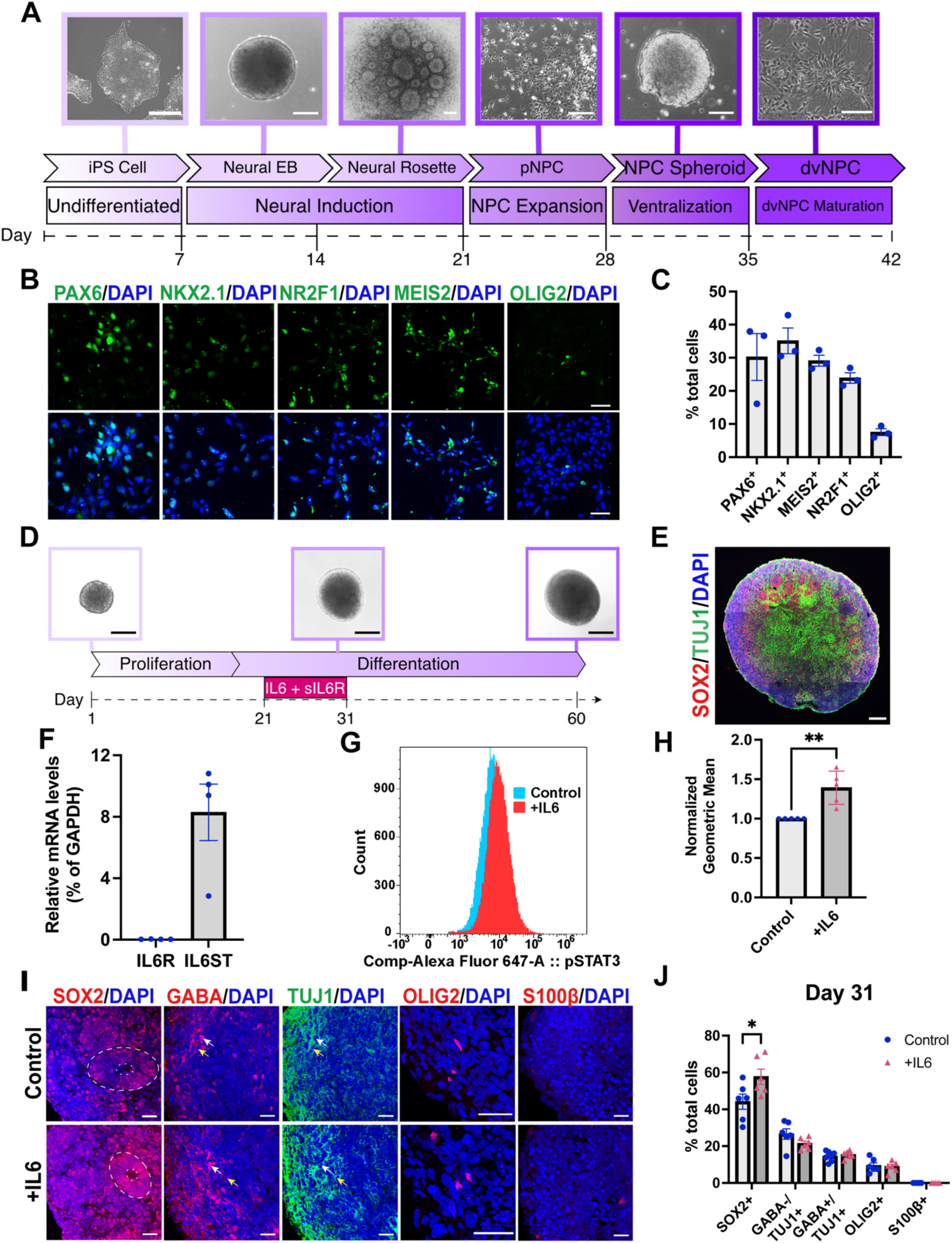
Dorsoventrally patterned forebrain organoids respond to IL-6 stimulation, resulting in expansion of the SOX2+ progenitor pool. (A) Schematic procedure for generating ventral neural progenitor cells (vNPCs) from human pluripotent stem cells. (B and C) Representative images (B) and quantification (C) of PAX6, NKX2.1, NR2F1, MEIS2, and OLIG2-expressing vNPCs following patterning with Pur (n = 3 batches). (D) A schematic procedure for forebrain organoid generation, culture paradigm, and IL-6 treatment duration. (E) Representative image of SOX2+/TUJ1+ rosette-like structures in Day 31 organoids. (F) qPCR analysis of IL6R and IL6ST expression in Day 21 organoids (n = 4 independent experiments). (G and H) Representative flow cytometry plot (G) and quantification (H) of phosphorylated STAT3 (pSTAT3) expression in control and IL-6-treated organoids at Day 31 (n = 4 independent experiments). Unpaired t-test. ** p < 0.01. (I and J) Representative images (I) and quantification (J) of SOX2, GABA, TUJ1, OLIG2, and S100ß-expressing cells in control and IL-6-treated organoids collected at Day 31 (n = 6 batches). Unpaired t-test. * p < 0.05. All values represent means, and all error bars represent SEM. Scale bars = 50 µM (A,C,D, and E) and 20 µM (I).

We next generated forebrain organoids from these mixed dorso-ventral NPCs (dvNPCs) using a controlled initial seeding density of 10,000 cells per organoid to minimize size variability (Figure 1D). Furthermore, to ensure experimental consistency and minimize batch-to-batch variation in the proportions of the different dorsal and ventral cell type populations across batches, all organoids were generated from dvNPCs within the first passage. These organoids exhibited organized neuroepithelial structures, including rosette-like regions comprised of SOX2+ progenitors, and following differentiation, these rosettes were surrounded by TUJ1+ immature neurons, consistent with early developmental architecture (Figure 1E and Supplementary Data Figure 1A).

We next assessed whether the neural cells within these organoids were competent to respond to IL-6. In the human fetal brain, IL-6 signaling in NPCs is mediated by the secretion of the soluble IL-6 receptor (sIL-6R) from non-cell-autonomous sources. sIL-6R binds extracellular IL-6 and subsequently interacts with the transmembrane IL-6 signal transducer (IL6ST) to activate JAK/STAT signaling^35–38^. Accordingly, the organoids used in this study had nearly undetectable levels of IL6R on Day 21, the day we began our IL-6 exposure (Figure 1F), and strongly expressed the IL6ST, which aligns with previous analyses of IL-6 receptor components in human pluripotent stem cell (hPSC)-derived NPCs^31,39,40^.

Previous studies also demonstrated that simultaneous treatment with IL-6 and sIL-6R stimulated JAK/STAT activation in hPSC-derived neural cells^31,39,40^. Therefore, to investigate the effects of IL-6 signaling, organoids were treated with 8.8 ng/mL (421.1 pM) of recombinant human IL-6 and an equimolar concentration of recombinant human sIL6R (16 ng/mL) for 10 days. The IL-6 concentration and treatment duration were selected to match those used in previous studies^31^. The sIL-6R concentration was selected to remain within physiologically relevant levels, as cerebrospinal fluid sIL-6R concentrations have been reported to remain within physiological ranges even under inflammatory conditions^41^. To confirm pathway activation, we assessed changes in phosphorylated STAT3 (pSTAT3), the core downstream effector of IL-6 signaling, using flow cytometry. By the end of the 10-day stimulation period, pSTAT3 was significantly elevated in the IL-6-treated organoids compared to untreated controls (Figure 1G, H). Together, these data confirm that co-treatment with recombinant IL-6 and sIL-6R is sufficient to robustly activate canonical IL-6 signaling in our organoids, validating this system as suitable to investigate the downstream effects of IL-6 exposure on neuronal development.

We next evaluated how IL-6 altered neural development using this new organoid platform. Organoids were stimulated daily with IL-6 and sIL-6R beginning at 21 days *in vitro* and then stained on Day 31 (Figure 1I). Notably, we observed a significant increase in the SOX2+ progenitor cell population in IL-6-treated organoids compared to untreated organoids, as measured by total percentage of SOX2+ nuclei. In contrast, the proportions of GABA+ interneurons and TUJ1+ neurons remained unchanged, indicating that early neuronal differentiation was not detectably affected at this stage. Similarly, the percentage of OLIG2+ progenitors remained unaltered, and S100β+ astroglia were not yet detectable, consistent with the early developmental stage of the organoids. These results suggest that IL-6 primarily acts on neural progenitors, potentially by modulating proliferation dynamics or progenitor state, rather than directly altering neuronal differentiation. This interpretation is in agreement with prior studies that identified radial glia as preferentially responsive to elevated levels of IL-6 during neurodevelopment^31^.

### IL-6 induces a neuroinflammatory transcriptional program coupled to early synaptic gene dysregulation

To capture early transcriptional responses to IL-6, we performed bulk RNA-sequencing (RNA-seq) on organoids after 10 days of IL-6 treatment. Two pairs of organoids, control and IL-6-treated, were collected from two different hiPSC lines for a total of eight samples (Supplementary Figure 1B). By comparing our dataset with a previously published transcriptomic dataset from the human fetal brain, we found that our samples were most similar to first-trimester tissue, positioning our model within an early neurodevelopmental window^42^ (Figure 2A). Differential gene expression analysis revealed a specific transcriptional change dominated by the upregulation of inflammatory signaling genes alongside dysregulation of genes involved in synaptic function (Figure 2B, Supplementary Table 1). Canonical IL-6 pathway targets, including STAT3 and A2M, were among the most significantly upregulated genes, confirming pathway activation at the transcriptional level. Additionally, increased CFI expression, which encodes a key negative regulator of complement activation, suggests active engagement of complement regulatory mechanisms in the IL-6-treated organoids. Interestingly, several interferon-associated genes were also elevated, including IRF9, B2M, and NLRC5^43^. NLRC5 drives major histocompatibility complex class I (MHC-I) expression in neurons, and its downstream target B2M has been implicated in synaptic remodeling^44–48^. Moreover, upregulation of MHC-I genes has been reported in other MIA studies and has also been associated with ASD^31,49–53^, potentially linking the inflammatory response to altered synaptic development.

**Figure 2.**
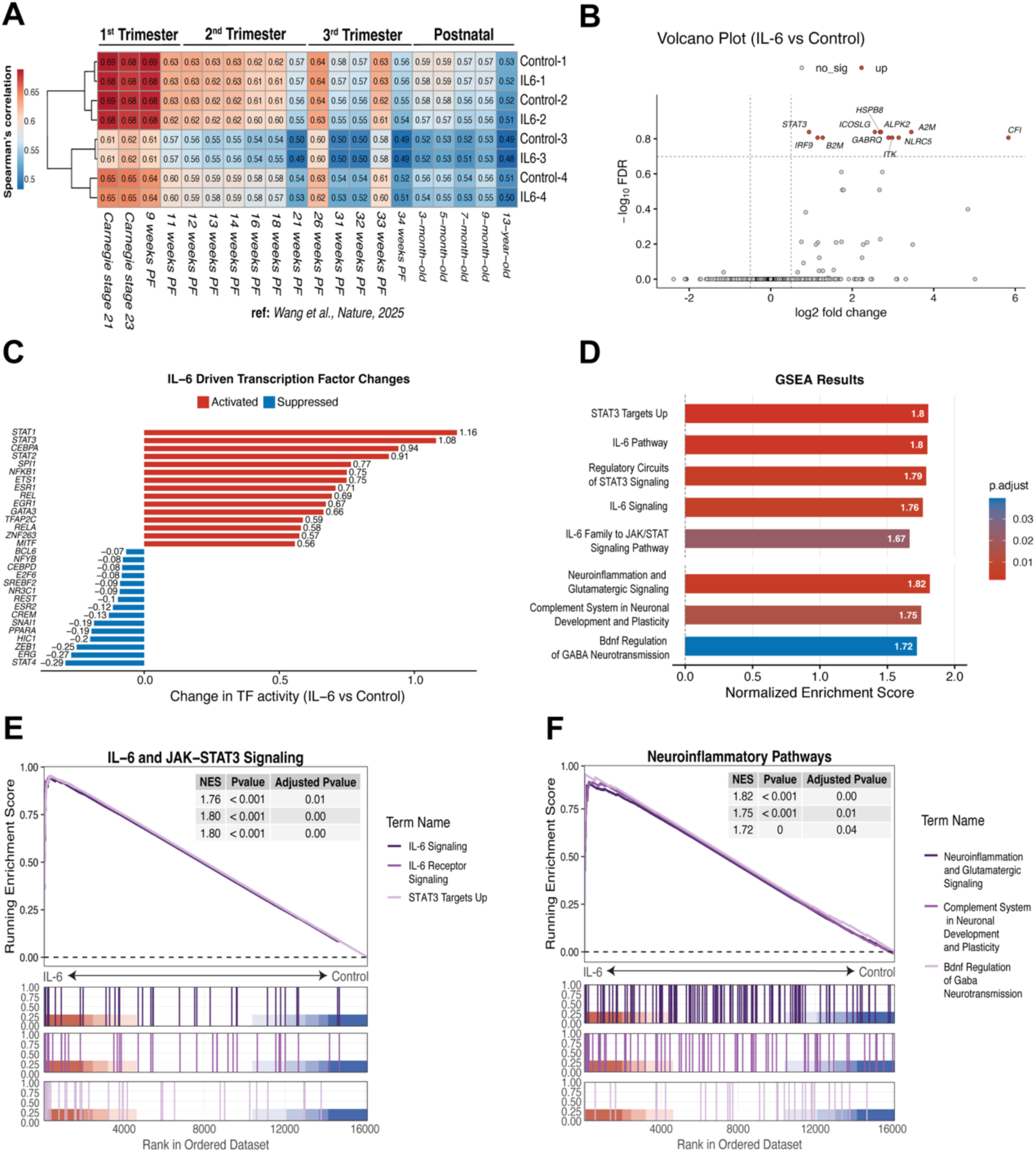
Bulk RNA-sequencing reveals inflammatory pathway activation and early inhibitory signaling disruption on Day 31. (A) Transcriptomic age correlation between control and IL-6-treated organoids and human fetal brain samples in reference dataset Wang et al., 2025^42^. (B) A volcano plot comparing differentially expressed genes in control and IL-6-treated organoids. Differential expression was evaluated using a Quasi-likelihood F-test implemented in edgeR. Multiple testing was corrected using the Benjamini-Hochberg method to control the false discovery rate (FDR). Significantly upregulated genes are in red (|log2 Fold Change| > 0.5 and FDR < 0.2). (C) Bar chart showing the change in transcription factor (TF) activity between IL-6-treated and control conditions, estimated using DoRothEA. TFs are ranked by magnitude of activity change (IL-6 vs control). Red bars indicate activated TFs; blue bars indicate suppressed TFs. (D) GSEA results showing normalized enrichment scores (NES) for selected gene sets from the MSigDB C2 curated gene set collection. Gene sets are grouped by categories: IL-6/JAK-STAT signaling (top) and neuroinflammatory pathways (bottom). Bar color reflects the Benjamini-Hochberg adjusted p-value, with darker red indicating greater significance. (E and F) GSEA enrichment plots for (E) IL-6 and JAK-STAT3 signaling gene sets and (F) neuroinflammatory pathway gene sets. The running enrichment score, gene hit positions, and rank metric distribution are shown. NES, nominal p-value, and Benjamini-Hochberg adjusted p-value are inset for each gene set.

Transcription factor analysis pointed to more specific mechanisms underlying this synaptic dysregulation. DoRothEA transcription factor analysis revealed enhanced activation of EGR1, a regulator of neural cell proliferation and synaptic plasticity^54,55^, and suppression of REST, a repressor of neuronal gene program activation in progenitor cells (Figure 2C, Supplementary Table 2). EGR1 enrichment has been found to be necessary to increase levels of EGFR in NPCs^55^, consistent with our observations of progenitor expansion (Figure 1I,J), suggesting a potential mechanism through which IL-6 drives this effect. REST suppression after IL-6 exposure suggests that neuroinflammation may permit a shift in progenitor state that promotes earlier neuronal commitment and differentiation^56^. Consistent with this hypothesis, we detected significant upregulation of GABRQ, which encodes the GABA-A receptor theta subunit, suggesting perturbations in GABA signaling (Figure 2A). Prior evidence showed that EGR1 directly binds to and transcriptionally activates GABRQ, providing a pathway through which IL-6-driven transcriptional changes can dysregulate GABAergic signaling^57^. Gene set enrichment analysis (GSEA) corroborated these findings, showing positive enrichment of IL-6/JAK-STAT pathway related terms, including “IL-6 Signaling”, “Regulatory Circuits of STAT3 Signaling”, and “STAT3 Targets Up”, as well as terms associated with neuronal function, including “Neuroinflammation and Glutamatergic Signaling” and “BDNF Regulation of GABA Neurotransmission” (Figure 2D-F).

Collectively, these findings suggest that transiently elevated IL-6 levels elicit an inflammatory response in prenatal neural lineage cells that affects both canonical inflammatory programs and genes implicated in GABA signaling, aligning with transcriptional signatures observed in MIA models of ASD and related neurodevelopmental conditions.

### Single-nucleus RNA-seq reveals cell-type-specific effects of IL-6

As the bulk RNAseq data showed that IL-6 stimulated an acute inflammatory response and that it promoted the expansion of the SOX2+ progenitor pool with modest changes in transcriptional programs underlying neuronal functional development, we next sought to establish the longer-term consequences of IL-6 exposure on neural development. When the cellular composition of the organoids was evaluated at Day 60,1 month after IL-6 treatment, there was a trend towards increased numbers of GABA+/TUJ1+ interneurons, whereas the GABA-/TUJ1+ presumptive glutamatergic neurons remained unchanged. Of note, we observed a significant increase in astroglial differentiation, as indicated by S100β acquisition. However, we no longer detected cells expressing OLIG2(Figure 3A,B).

**Figure 3.**
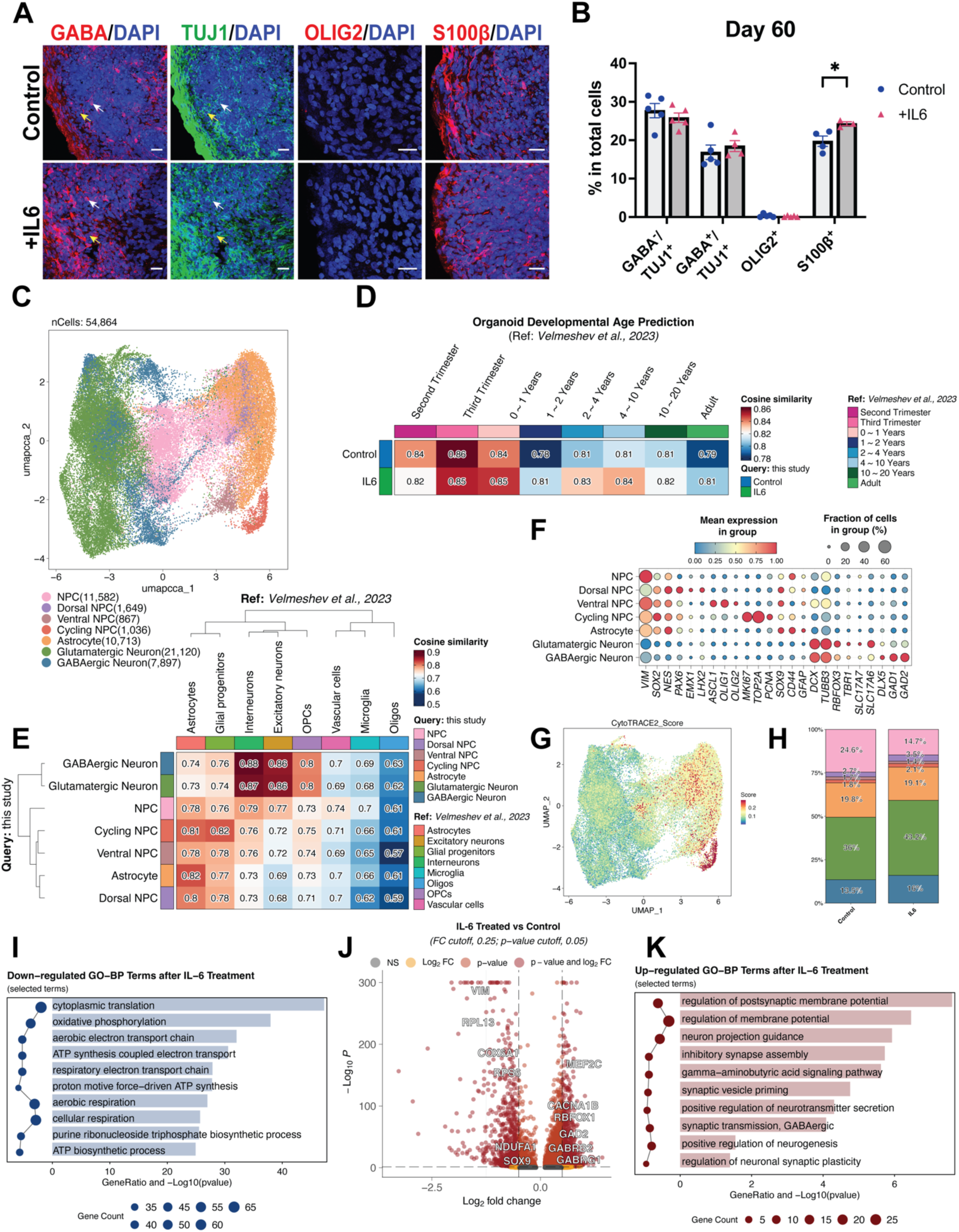
Single-nucleus RNA sequencing uncovers cell-type-specific effects on neuronal maturation on Day 60. (A and B) Representative images (A) and quantification (B) of GABA, TUJ1, OLIG2, and S100ß-expressing cells in control and IL-6-treated organoids at Day 60 (n = 5 batches). All values represent means, and all error bars represent SEM. Scale bar = 20 µM. Unpaired t-test. * p < 0.05 for each gene set. (C) Uniform manifold approximation and projection (UMAP) plot showing neural lineage clusters from control and IL-6-treated organoids. (D) Heatmap indicating the age similarity between Velmeshev et al., 2023^58^ and this study. (E) Heatmap showing the cell-type similarity between Velmeshev et al., 2023^58^ and this study. (F) Dot plot demonstrating the expression of neural lineage cell marker genes of each annotated cell type. (G) UMAP plot displaying CytoTrace2 differentiation state scores. Red indicates less differentiated; blue indicates more terminally differentiated cells. (H) Proportion plot representing the percentage of each cell type in control vs IL-6-treated organoids. (I) Gene ontology (GO) Biological Process (BP) enrichment analysis of downregulated differentially expressed genes (DEGs) in control vs IL-6-treated organoids. (J) Volcano plot showing the DEGs between control and IL-6-treated organoids (K) Gene ontology (GO) Biological Process (BP) enrichment analysis of downregulated differentially expressed genes (DEGs) in IL-6-treated vs control organoids.

To further examine sustained molecular changes in neuronal maturation, we performed single-nucleus RNA sequencing (snRNA-seq) one month after IL-6 treatment. Using two batches of control and IL-6-treated organoids generated from two different cell lines, we obtained a total of 54,864 nuclei with a median of 1,351 unique transcripts per nucleus after quality control (Supplementary Figure 1C,D). Shared nearest neighbor (SNN) clustering identified 17 clusters representing 7 major cell types (Figure 3C and Supplementary Figure 1E). Correlation analysis with an existing human developing cortical tissue snRNA-seq dataset showed that the organoids at this time point most resembled the third-trimester fetal brain rather than the second trimester or postnatal periods of development^58^ (Figure 3D). Comparison with this same dataset validated the neuronal and glial identities of maturing cells in each cluster^58^ (Figure 3E). Moreover, the cells in these clusters expressed canonical markers indicative of progenitor cells (SOX2, NES, VIM), glutamatergic neurons (DCX, TUBB3, SLC17A6), GABAergic interneurons (GAD1, GAD2, DLX5), and astrocytes (SOX9, CD44, GFAP) (Figure 3F). The progenitor cell populations were subdivided into Dorsal NPCs, (PAX6, EMX1, LHX2), Ventral NPCs (ASCL1, OLIG1, OLIG2), and Cycling NPCs (MKI67, TOP2A, PCNA) (Figure 3D). These developmental states were further validated using CytoTRACE, which showed decreasing differentiation potential from the cycling NPCs through to the differentiated glutamatergic and GABAergic neuron populations (Figure 3G).

After identifying the transcriptomic profiles of the cell types comprising the organoids at this time point, we next determined whether there were changes in the cell type composition of organoids that had been transiently exposed to IL-6. In contrast to organoids analyzed at the end of IL-6 exposure on Day 31, this analysis found a significant decrease in NPCs, consistent with a model in which transient IL-6 first promotes progenitor proliferation before accelerating depletion of the stem cell pool (Figure 3H and Supplementary Data Figure 1F,G). This is similar to what has been previously observed in mouse studies, where IL-6 exposure in the perinatal period promoted progenitor proliferation before exhausting the stem cell pool more quickly than in mice that did not receive a transient increase in peripheral IL-6^59^. This reduction in NPCs was accompanied by increases in both glutamatergic and GABAergic neuron populations, suggesting that the early progenitor expansion may have ultimately produced a larger neuronal output. Furthermore, in agreement with this observation, our differential gene expression analysis comparing IL-6-treated to control cells also revealed upregulation of genes related to interneuron maturation, such as GAD2, the GABA receptor unit-encoding genes GABRB2 and GABRG1, and RBFOX1, which has a known role in supporting interneuron maturation and synaptic function (Figure 3J, Supplementary Table 3) ^60,61^. Gene ontology analysis indicated that biological processes associated with maturing interneuron function, such as inhibitory synapse assembly, the GABA signaling pathway, and GABAergic synaptic transmission, were enriched, suggesting accelerated activation of inhibitory synaptic machinery, likely within the GABAergic neuronal cluster (Figure 3K, Supplementary Table 4). This enrichment of a GABAergic neurotransmission signature is notable, as our bulk RNA-seq analysis at Day 31 had already hinted at potential disruptions in GABAergic neuron development. However, the simultaneous downregulation of processes such as cytoplasmic translation and oxidative phosphorylation, and related metabolic processes, raises the possibility that premature activation of maturation programs may exceed the metabolic capacity required to sustain them (Figure 3I, Supplementary Table 5). Altogether, these findings suggest that IL-6 drives premature activation of interneuron maturation programs, pushing cells toward transcriptional states that surpass their expected developmental stage.

### Transient levels of IL-6 impair long-term interneuron maturation

Since we observed marked changes in interneuron maturation programs, we next focused on the interneuron lineage cells in our dataset. To this end, we re-clustered the NPCs, Ventral NPCs, Cycling NPCs, and GABAergic Neuron clusters (Figure 4A and Supplementary Data Figure 1H). This analysis enabled us to characterize cells as Interneuron Progenitors, Immature Interneurons, and Mature Interneurons (Figure 4B). Interneuron Progenitors exhibited enriched expression of PVALB, NPY, and LHX6, all markers of the MGE lineage with minimal expression of immature and mature neuronal markers (Supplementary Figure 1I). This expression pattern, particularly PVALB, which is typically upregulated in mature interneurons, may indicate MGE lineage-fate priming rather than terminal fate commitment^30,62^. Immature Interneurons expressed early neuronal markers such as TUBB3, DCX, and STMN2, whereas Mature Interneurons expressed RBFOX3, GAD1, and GAD2.

**Figure 4.**
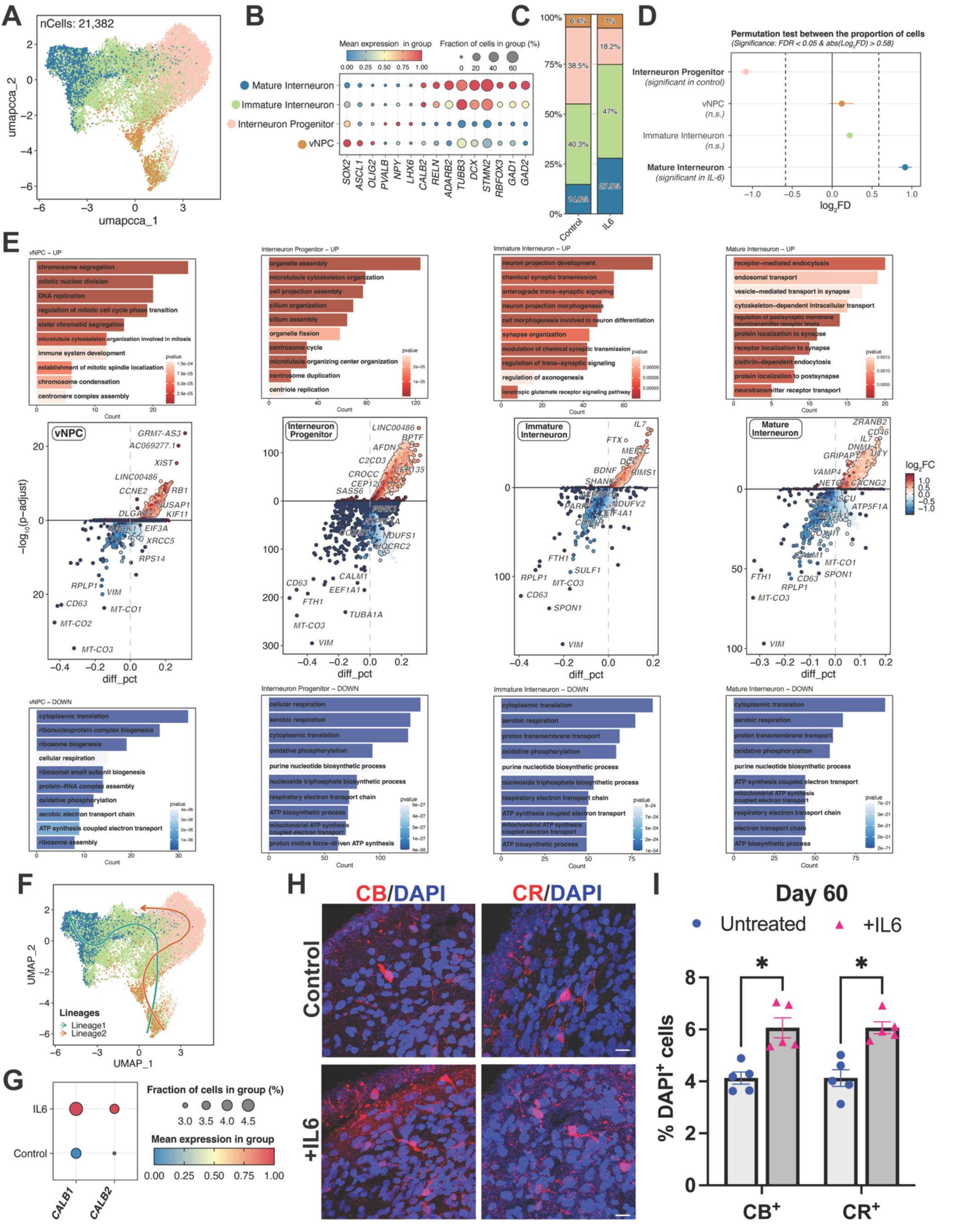
Single-nucleus RNA sequencing uncovers cell-type-specific effects on neuronal maturation on Day 60. (A) Uniform manifold approximation and projection (UMAP) plot showing interneuron lineage clusters from control and IL-6-treated organoids. (B) Dot plot indicating the expression of interneuron lineage cell marker genes of each annotated cell type. (C) Proportion plot representing the percentage of each cell type in control vs IL-6-treated organoids. (D) Permutation plot demonstrating the fold change and significance per interneuron lineage cluster between control and IL-6-treated organoids. (E) DEG and pathway enrichment analysis across interneuron lineage stages in IL-6 vs control conditions. (*Top*) Bar plots depicting significantly enriched GO BP terms among upregulated genes in each population. (*Middle*) Volcano plots displaying DEGs. (*Bottom*) GO BP terms enriched among downregulated genes in each population. (F) UMAP visualization displaying the inferred pseudotime trajectories (G) Dot plot demonstrating the CALB1 and CALB2 expression levels in control vs IL-6-treated organoids. (H and I) Representative images (H) and quantification (I) of calbindin (CB) and calretinin (CR)-expressing cells in control vs IL-6-treated organoids (n = 5 batches). Scale bar =20 µM. Unpaired t-test. * p < 0.05.

Within the Immature and Mature Interneuron clusters, there were also cells expressing CALB2, RELN, and ADARB2, which are all enriched in CGE-derived interneurons. The IL-6-treated organoids have significantly fewer Interneuron Progenitors and significantly more Mature Interneurons than the controls, suggesting that transient IL-6 exposure accelerates progression through the interneuron lineage, depleting the progenitor pool while driving cells toward terminal differentiation (Figure 4C,D).

To determine where along the interneuron trajectory IL-6-induced changes occurred, we performed differential gene expression and gene ontology analyses comparing control and IL-6-treated cells across all four interneuron lineage clusters (Supplementary Tables 6-17). Broadly, transcriptional states across the interneuron lineage further demonstrate enhanced maturation, along with suppression of cytoplasmic translation and metabolic programs (Figure 4E). Strikingly, within the Ventral NPC cluster, Immune System Development was an enriched term, suggesting that these cells retain an inflammatory-like transcriptional state at least one month after IL-6 withdrawal. This may represent a neuroinflammatory state contributing to the altered maturation observed across the interneuron lineage. Other enriched terms in the Ventral NPC cluster, such as Chromosome Segregation and DNA Repair, are associated with the transition from progenitor cell to differentiated neuron. The Interneuron Progenitor cluster showed enrichment for terms related to structure and cytoskeletal reorganization, including Microtubule Cytoskeletal Organization, Cell Projection Assembly, and Neuron Projection Morphogenesis, processes consistent with early morphological changes during neuronal commitment. Within the Immature Interneuron cluster, we began to observe an upregulation of terms related to neuronal and synaptic maturation, including Neuron Projection Development, Synapse Organization, and Regulation of Trans-synaptic Signaling, indicating that interneurons in IL-6-treated organoids may be acquiring synaptic properties more rapidly. Similarly, in Mature Interneurons, upregulated terms such as Receptor-Mediated Endocytosis, Protein Localization to Synapse, and Neurotransmitter Receptor Transport suggest these cells may also be more functionally active. Common to all four clusters was the downregulation of Cytoplasmic Translation, Aerobic Respiration, and other metabolic processes, potentially representing long-term alterations in metabolic capacity following IL-6 exposure. Together, these data support the idea that changes in the interneuron lineage drive the observed global developmental changes seen in Figure 3. Moreover, they demonstrate a potential link between persistent inflammatory effects in the Ventral NPCs, which may be affecting the number of interneurons or, alternatively, the functional maturation of later-stage interneurons.

Given the transcriptional evidence for accelerated synaptic maturation and increased interneuron abundance, we next asked whether interneuron subtype specification was similarly affected. Interestingly, pseudotime analysis inferred two distinct lineages within the interneuron clusters (Figure 4F). The first lineage extended through the Ventral NPCs, Immature Interneuron, and Mature Interneuron clusters (Supplementary Data Figure 1J). The second lineage spanned the Ventral NPCs through Interneuron Progenitor clusters. Together with the subtype-specific marker expression patterns observed in Interneuron Progenitors and Mature Interneurons, this suggests that these two lineages may recapitulate divergent MGE-like and CGE-like fates. Importantly, this finding further establishes the validity of our ventralization paradigm in capturing diverse interneuron origins. GABAergic interneurons typically begin to express subtype-specific markers later in development. Therefore, we assessed the production of calbindin (CALB1^+^) and calretinin (CALB2^+^), markers for two interneuron subpopulations predominantly derived from the CGE. From our snRNA-seq analysis, we detected increased expression of both CALB1 and CALB2 at the transcriptome level (Figure 4G). Immunohistochemistry similarly demonstrated a significant increase in CB^+^ and CR^+^ cells (Figure 4H,I). This enhancement in CB^+^ and CR^+^ production is consistent with accelerated neuronal differentiation following IL-6 exposure. However, given the concurrent reduction in MGE-like progenitor proportions, a complementary interpretation is that IL-6 biases fate specification toward CGE-derived interneuron subtypes at the expense of MGE-derived ones.

### IL-6 exposure persistently alters gene regulatory networks associated with inflammation within the interneuron lineage

To interrogate gene regulatory networks (GRNs) altered by transient IL-6 exposure, we next performed *de novo* transcription factor enrichment and regulon inference using pySCENIC. This revealed 134 regulons enriched in our dataset. These regulons showed differential activity across the four interneuron lineage clusters and between control and IL-6-treated cells (Figure 5A, Supplementary Table 18). We then used a connection specificity index to identify regulon modules based on pairwise co-activation (Figure 5B, Supplementary Table 19). This analysis assigned the 134 regulons into four discrete modules. The regulons comprising Module 1 were primarily related to progenitor identity, including NANOG, POU5F1B, and TCF4, as well as those that regulate the cell cycle, such as E2F2, E2F8, and MYBL1. Other regulons indicated early fate priming in these cells: FOXN4, THRB, TFAP2C, and ETV5, together indicating this to be a transitional progenitor state. Module 2 was defined by regulons that repress proliferation (RB1, MXI1) and by epigenetic modifications that facilitate cell-fate commitment (CHD1/2, ARID3A). This coincided with the co-activation of transcriptional programs related to neural lineage specification (NKX2-2, FOXP1/2). Module 2 also included MECP2, whose role in interneuron maturation and GABAergic circuit development is well established, and whose loss-of-function causes Rett syndrome^63,64^. The correlated activation of regulons in Module 3, such as SOX4, SOX11, POU2F1/2, and BCL11A, corresponds to the terminal differentiation of neurons. Interestingly, we also detected a correlated subset suggestive of stress and immune responsiveness, including STAT1, ATF1, BACH2, and ING4. The appearance of this signature is consistent with a sustained inflammatory transcriptional state following transient IL-6 exposure.

**Figure 5.**
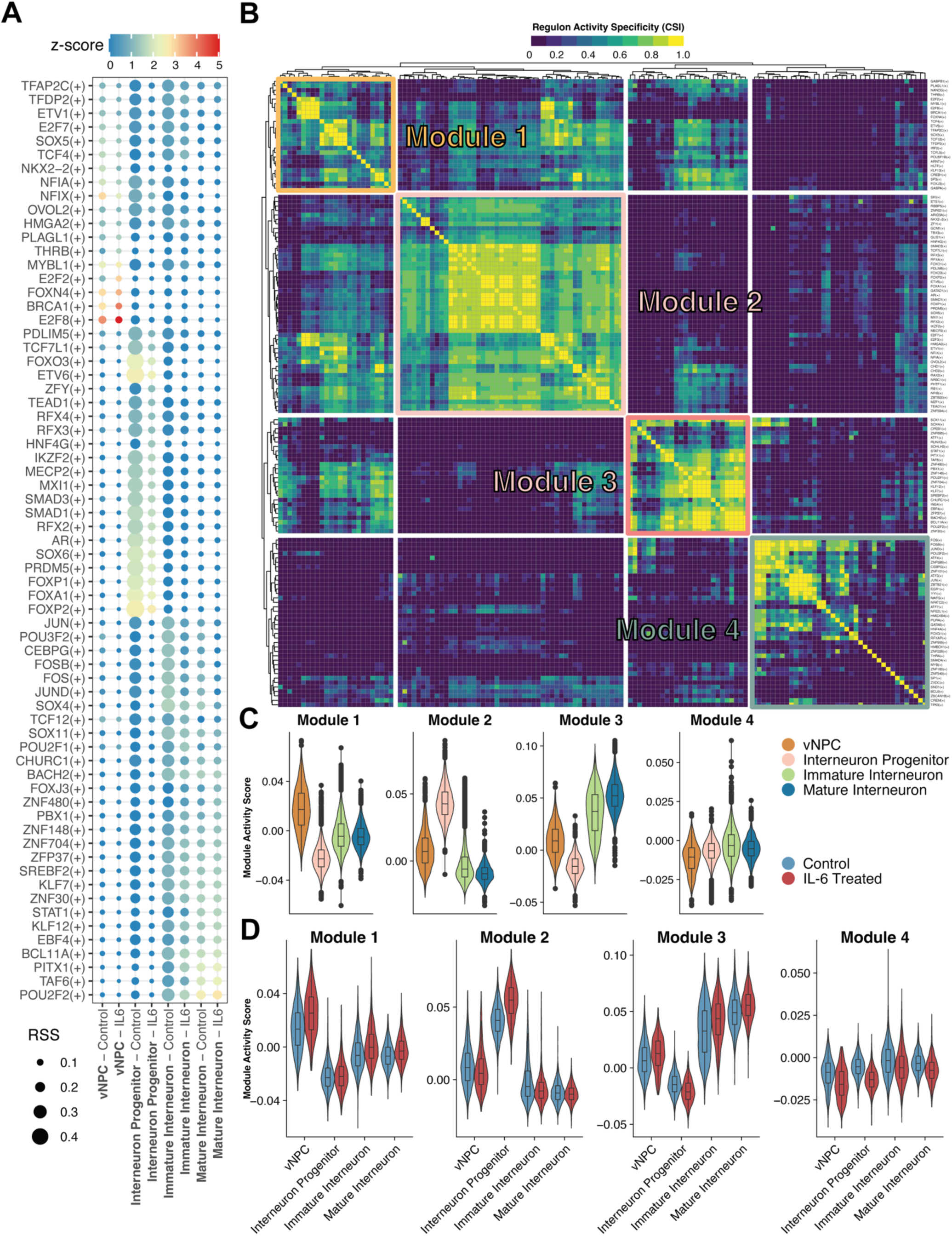
*De novo* regulon and gene regulatory network analysis of interneuron-lineage cells. (A) Regulon specificity score (RSS) analysis showing cell type-specific regulon activity across annotated interneuron-lineage cell types under control and IL-6-treated conditions. (B) Heatmap showing regulon modules identified by hierarchical clustering of the Connection Specificity Index (CSI) matrix. (C) Violin plots showing regulon module activity scores across annotated interneuron-lineage cell types. (D) Violin plots comparing regulon module activity scores between control and IL-6-treated conditions within annotated interneuron-lineage cell types.

Enhanced activation of Modules 1-3 was developmentally regulated (Figure 5C). For instance, Module 1 had the highest activity score in the Ventral Progenitor cluster, Module 2 had a greater activity score in the Interneuron Progenitor cluster, and Module 3 was most enriched in the Immature Interneuron cluster. Additionally, Module 3 expression appears to stabilize in the Mature Interneuron cluster, corresponding with the attainment of greater neuronal maturation. Compared with control cells, Module 1 shows greater activation in IL-6-treated cells within the Ventral Progenitor cluster (Figure 5D). This is striking, given the overall decrease in the proportion of Ventral Progenitor cells. However, this may reflect maintenance of self-renewal properties even as other cells begin to differentiate. This agrees with what we observe in the Module 2 activity, where IL-6-treated Interneuron Progenitors have a higher activity score than control Interneuron Progenitors, suggesting that a greater proportion of IL-6-treated cells are upregulating transcriptional programs that enable neural fate commitment. Module 3 programs similarly trend toward greater enrichment in IL-6-treated Immature and Mature Interneurons compared with controls, which parallels the increase in Mature Interneurons we observed in our cell type abundance analysis.

In contrast to Modules 1-3, Module 4 was characterized entirely by the activation of programs related to stress response, inflammatory signaling, and damage repair. This includes broad activation of the AP-1 transcription factor family, as well as EGR1, the early response gene that was highly activated immediately following IL-6 stimulation (Figure 5B). The activity score for Module 4 was relatively stable across all four clusters, suggesting that this inflammatory signature is conserved across developmental states rather than restricted to progenitors (Figure 5C). Notably, however, Module 4 activity was higher in control cells than in IL-6-treated cells at Day 60 (Figure 5D). The presence of this signature in control cells indicates that a baseline level of stress-responsive transcription is typical in this system, particularly in interneuron development, rather than an artifact of IL-6 treatment. Its attenuation in IL-6 cells could reflect either a suppression of immune response programs following the initial cytokine stimulus, or a blunting of inflammatory responsiveness, demonstrating a tolerance to inflammatory stimulation following acute exposure. A similar dampening was observed in a mouse model of MIA, in which microglia in the developing brain exhibited reduced responsiveness to inflammatory stimuli following prenatal stress that persisted throughout development^65^. However, whether a similar mechanism occurs in human neural lineage cells remains to be determined. Together, the progressive enhancement of developmental programs in IL-6-treated cells, along with diminished inflammatory reactivity in mature interneurons, suggest that transient IL-6 exposure leaves a persistent and detectable transcriptional mark. These findings have potential implications for neurodevelopmental consequences of prenatal immune activation on dysregulated interneuron maturation, particularly in ASD^66,67^.

## Discussion

The present study demonstrates that transient elevation of IL-6 during a critical window of human forebrain organoid development produces lasting transcriptional changes within the interneuron lineage, with evidence of accelerated maturation programs, altered subtype specification, and a persistent inflammatory transcriptional state after IL-6 withdrawal. These findings extend previous studies of MIA by providing detailed insights into how prenatal IL-6 exposure disturbs human interneuron development.

Mounting evidence points to disruption in interneuron development as a downstream consequence of MIA, particularly in rodent models that converge on interneuron deficits, following *in utero* inflammation^15–17,68–70^. However, the translational relevance of these findings to humans has remained unanswered, in part because mouse and human brain development are temporally incongruent^25,71^. In rodents, much of interneuron development occurs during mid-gestation^72^, whereas in humans, interneurons continue to emerge and migrate throughout the second and third trimesters, with maturation extending well into the postnatal period. Therefore, the timing of immune activation may differentially affect interneuron development and maturation in humans compared to mice. Consistent with this, one study using a murine model demonstrated that the timing of inflammatory stimulus exposure had cell-type-specific effects on interneuron subtype output^69^, suggesting that the developmental stage at which immune activation occurs is highly instrumental in the cellular outcomes. Our hiPSC-based model addresses this translational gap through its transcriptional correspondence with defined stages of human fetal development. Transcriptomic age analysis confirmed that organoids at the time of IL-6 exposure most closely resembled the first trimester human fetal brain, a period when interneuron fate remains plastic and ventral forebrain identity consolidation is actively underway^25,30,73^. By Day 60, the organoids had advanced to a transcriptional state most similar to third-trimester tissue, a period characterized by interneuron maturation and subtype specification^74–76^. This developmental progression allowed us to model both the initial response and its downstream consequences within biologically relevant stages.

Using our model system, we provide evidence that IL-6 exposure during the prenatal window is sufficient to alter interneuron maturation trajectories beyond the period of IL-6 exposure. Several findings from our model are in agreement with previous reports in rodent models, hPSC-based systems, and human tissue, supporting its validity for studying MIA. The expansion of SOX2+ progenitors aligns with observations from both prior organoid studies^31^ and mouse models of *in utero* IL-6 exposure^59,77,78^. Additionally, the transcriptional upregulation of MHC-I pathway components recapitulates signatures observed in other MIA models and in postmortem ASD brain tissue^31,53,79,80^. Interestingly, the IL-6 family cytokine LIF was also observed to increase the production of CGE-like interneurons in a hPSC-derived organoid model, providing further evidence that IL-6 family cytokines can promote an interneuron fate^81^.

Our observed increase in mature interneurons at Day 60 likely reflects both enhanced interneuron production and accelerated maturation. The reduction in interneuron progenitors suggests that cells are progressing more rapidly through early stages rather than accumulating at mature states. Consistent with this, IL-6–treated cells precociously expressed maturation-associated transcriptional programs—including GAD2, GABRB2, GABRG1, and RBFOX1—along with enrichment of gene ontology terms related to inhibitory synapse assembly and GABAergic transmission, supporting premature activation of late-stage functional programs. These findings also align with a previous study using hPSC-derived brain organoid models generated from idiopathic ASD lines, which observed enhanced interneuron output and synapse abundance, suggesting these may represent convergent biology underlying ASD regardless of genetic background^82^. In parallel, transcriptional evidence of accelerated maturation was accompanied by altered interneuron subtype specification. The increase in CALB1⁺ and CALB2⁺ interneurons, together with a reduction in MGE-like progenitor populations, suggests that IL-6 biases fate specification toward CGE-derived interneuron subtypes at the expense of MGE-derived lineages. This observation aligns with reported reductions in parvalbumin-positive interneurons in the prefrontal cortex of individuals with ASD^83^, and across multiple MIA studies in rodent models and hPSC-derived systems^15–17,24,68–70^. Notably, a similar shift in MGE to CGE transcriptional programs was observed following loss-of-function of two ASD-associated genes in an hPSC-derived assembloid model, suggesting that this may also be a convergent vulnerability across both genetic and environmental ASD risk factors^84^. Supporting this, our regulon analysis using pySCENIC identified Module 2 as enriched for transcription factors associated with interneuron maturation and MGE identity, including SOX6^85^ and NFIB^86^, whereas Module 3 contains transcription factors linked to terminal neuronal differentiation programs that have been identified in CGE-enriched interneuron lineages in prior organoid-based regulon analyses, including SOX4 and SOX11^87^. Moreover, MGE-derived interneurons are known to have a prolonged maturation timeline extending into postnatal development in humans^75,88^, and the MGE-like progenitor population appears to be immature relative to more fate-committed CGE-like interneurons in our system. Whether these modules reflect distinct MGE- and CGE-like transcriptional programs differentially engaged by IL-6 remains to be determined. This distinction is biologically important, as MGE-derived parvalbumin-positive interneurons play a central role in regulating network excitability and synchronizing cortical oscillations via fast inhibition of pyramidal neurons^89^, and CGE-derived interneurons provide more indirect inhibitory control by inhibiting other interneurons^90^. Interestingly, stereological counts of interneuron subpopulations in sections from ASD individuals revealed an increased number of calretinin+ interneurons in the hippocampus^91^. Elucidating these lineage-specific effects will have clear translational relevance, as disruption of excitation–inhibition balance is a well-established mechanism underlying core features of ASD.

The persistence of an inflammatory-like transcriptional state in Ventral NPCs one month after IL-6 withdrawal, as evidenced by the enrichment of Immune System Development GO terms and the sustained activity of stress- and immune-associated regulons in Module 4 from our regulon analysis, suggests that transient cytokine exposure leaves a lasting molecular mark on these progenitors. One possibility is that acute IL-6 stimulus induces epigenetic modifications, as previous studies have shown that JAK/STAT activation promotes DNA methylation changes that inhibit neurogenesis^92^, which would also be consistent with the concurrent accelerated maturation we observed at Day 60 in our organoid model. Notably, this pattern of sustained neuroinflammation may not be unique to the prenatal window, as IL-6 levels measured in postmortem forebrain tissue from ASD-diagnosed individuals through age 37 were approximately two-fold higher than in controls^93^, suggesting that the inflammatory state initiated prenatally may persist well into adulthood. This presents the possibility that prenatal IL-6 exposure establishes a neuroinflammatory state that may predispose the brain to increased risk of neurodegenerative disorders in adulthood. Supporting this possibility, several of the neuroimmune pathways activated in our system, including MHC-I upregulation driven by NLRC5/B2M and interferon-associated gene expression, have established roles not only in neurodevelopmental contexts but also in neurodegeneration. Complement-mediated synaptic pruning, MHC-I-dependent synaptic remodeling, and interferon-driven neurodegeneration have been recognized as contributors to neuronal loss and circuit disruption in aging and disease^94^. The activation of these same pathways during interneuron development suggests that the molecular machinery engaged by prenatal immune activation may prime neural circuits for later vulnerability. This is consistent with emerging epidemiological evidence that individuals with ASD have substantially elevated risks of developing dementia and Parkinson’s disease compared to the general population^95–98^, and this may reflect a shared neuroinflammatory mechanism linking early neurodevelopmental disruption to later degenerative processes.

The present study provides human cellular evidence that prenatal IL-6 exposure directly alters interneuron development and maturation. The persistent changes in gene expression described here also may perturb early synaptogenesis and lead to lasting defects in circuit assembly. Future studies using model systems that support human neural circuit formation, such as assembloid approaches^84,99,100^ or *in vivo* xenotransplantation of organoid-derived interneurons into chimeric brain models^26,101^, will be essential to define the developmental trajectory, long-term stability, and functional impact of IL-6–exposed interneurons. It is also important to note that our current organoid model lacks microglia, the brain’s resident immune cells. Given their responsiveness to IL-6 and established roles in complement-mediated synaptic refinement, incorporating microglia into these systems^102^ will be critical to determine whether the observed neuroimmune effects are amplified and further modulated by microglial activity. Finally, future work examining chromatin accessibility, epigenetic regulation, and the long-term stability of interneuron subtype composition will be necessary to determine whether these transcriptional changes translate into persistent circuit dysfunction relevant to neurodevelopmental disorders such as ASD.

## Acknowledgments

This work was supported in part by grants from the NIH (R21HD113311 to P.J. and S.W.L.) and the New Jersey Department of Health (CAUT24BRP009 to P.J. and S.W.L.). A.V.P. was supported by a T32 graduate training fellowship through the Training in Translating Neuroscience to Therapies program at Rutgers University (T32NS115700). M.J. was supported by a postdoctoral fellowship from the New Jersey Department of Health (CAUT24DFP004).

## Author Contributions

A.V.P. and P.J. designed the study, developed the experimental strategy, and interpreted the data. A.V.P. performed the majority of the experiments with technical assistance from M.N. and M.J. A.V.P. and Z.M. conducted RNA-seq analyses and interpreted sequencing datasets. S.W.L. provided critical input and guidance throughout the study. P.J. conceived and supervised the project. A.V.P. and P.J. wrote the manuscript with contributions and feedback from all co-authors.

---

**Figure S1.**
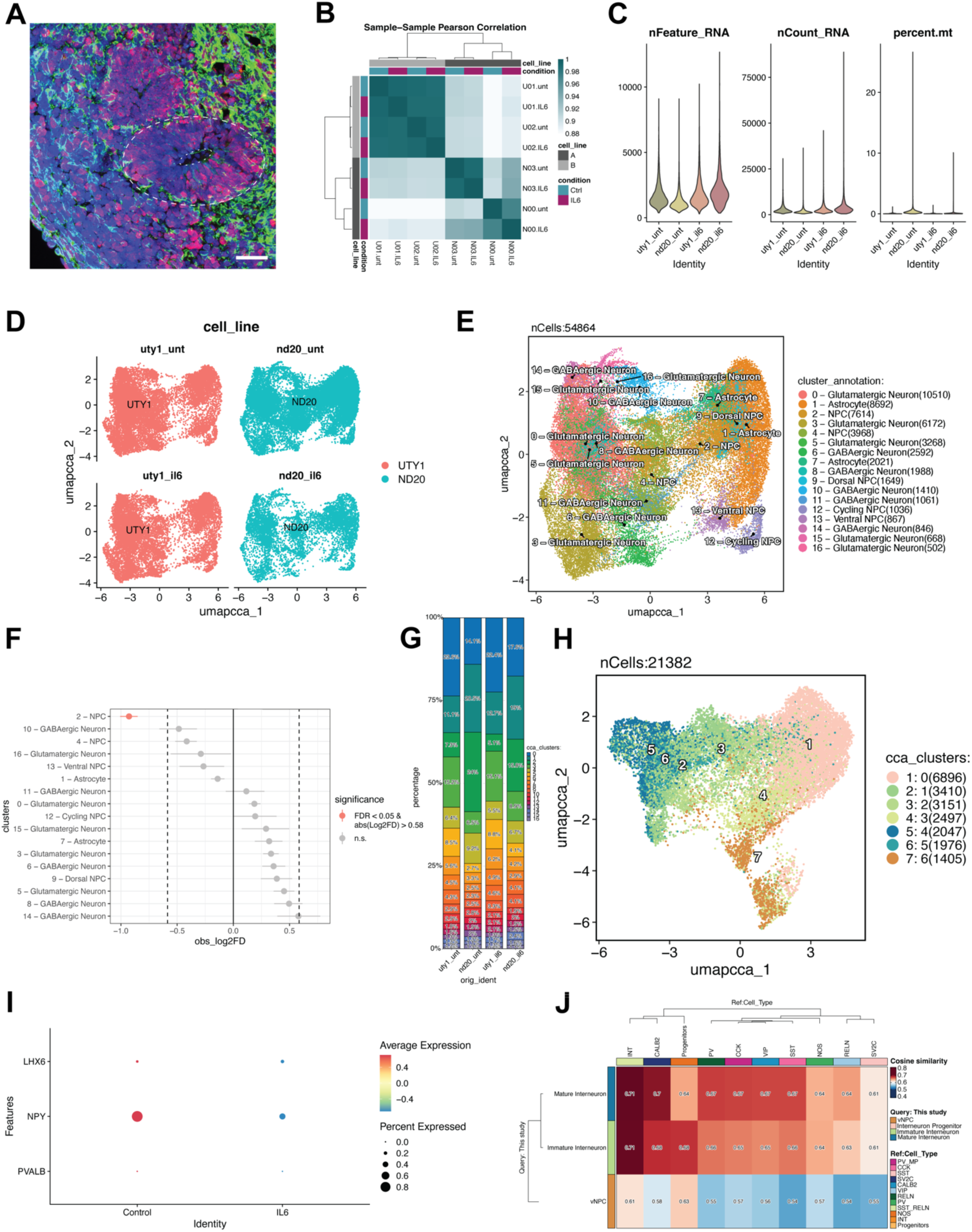
Quality control, integration, and annotation of the single-nucleus RNA-seq dataset (A) Representative image of a SOX2+/TUJ1+ rosette-like structure in Day 31 organoids. Scale bar = 20 µM. (B) Heatmap showing sample-sample correlation between bulk RNA-seq samples (n = 8; r = 0.88–0.98) (C) Violin plots showing quality control metrics per sample, including number of features (nFeature_RNA), number of counts (nCount_RNA), and mitochondrial read percentage (percent.mt), for each cell line and condition combination. (D) UMAP visualization of all cells colored by cell line (UTY1, ND20) and split by condition (Control, IL-6), demonstrating integration across batches. (E) UMAP visualization of all cell clusters following CCA integration. (F) Permutation plot presenting the significance of the proportion changes, grouped by cell type annotation, observed in Figure 3H. (G) Cell proportion plot for each cell line and condition across all clusters. (H) UMAP visualization of interneuron lineage cell subset clusters after CCA integration. (I) Dot plot showing average expression and percent of interneuron lineage cells expressing MGE-associated markers PVALB, LHX6, and NPY across control and IL-6 conditions. (J) Cosine similarity heatmap comparing interneuron lineage subclusters from pseudotime trajectory “Lineage 1” of this study (query) to cell types from the Velmeshev et al., 2023^58^ human fetal brain transcriptomic reference dataset.

